# Deficiency of the RNA-binding protein Cth2 extends yeast replicative lifespan by alleviating its repressive effects on mitochondrial function

**DOI:** 10.1101/2022.02.25.480133

**Authors:** Praveen K. Patnaik, Carine Beaupere, Hanna Barlit, Antonia María Romero, Mitsuhiro Tsuchiya, Michael Muir, María Teresa Martínez-Pastor, Sergi Puig, Matt Kaeberlein, Vyacheslav M. Labunskyy

## Abstract

Iron dyshomeostasis contributes to aging, but little information is available about the molecular mechanisms. Here, we provide evidence that, in *Saccharomyces cerevisiae*, aging is associated with altered expression of genes involved in iron homeostasis. We further demonstrate that defects in the conserved mRNA-binding protein Cth2, which controls stability and translation of mRNAs encoding iron-containing proteins, increase lifespan by alleviating its repressive effects on mitochondrial function. Mutation of the conserved cysteine residue in Cth2 that inhibits its RNA-binding activity is sufficient to confer longevity, whereas Cth2 gain-of-function shortens replicative lifespan. Consistent with its function in RNA degradation, we demonstrate that Cth2 deficiency relieves Cth2-mediated post-transcriptional repression of nuclear-encoded components of the electron transport chain. Our findings uncover a major role of the RNA-binding protein Cth2 in the regulation of lifespan and suggest that modulation of iron starvation signaling can serve as a target for potential aging interventions.

## Introduction

Iron (Fe) is an essential trace element, which serves as a cofactor for enzymes involved in multiple metabolic pathways, including synthesis of DNA, ribosome biogenesis, lipid metabolism, and mitochondrial oxidative phosphorylation. Dysregulation of Fe homeostasis has been implicated in the development of many human age-related diseases including diabetes (Simcox and McClain, 2013), inflammatory diseases (Verma and Cherayil, 2017), and neurodegeneration (Apostolakis and Kypraiou, 2017; Belaidi and Bush, 2016; Jiang et al., 2017; Stockwell et al., 2017), but little is known about the molecular details of how Fe homeostasis affects aging.

In eukaryotes, Fe utilization and storage are tightly controlled. Yeast *S. cerevisiae* contains two partially redundant transcription factors, Aft1 and Aft2, which activate the expression of genes known as Fe regulon that facilitate mobilization of intracellular Fe stores and its extracellular uptake in response to Fe deficiency (Kaplan and Kaplan, 2009; Philpott and Protchenko, 2008). In addition, Aft1 and Aft2 activate the expression of the RNA-binding protein Cth2. This protein contains two tandem zinc fingers (TZFs) that are required for binding AU-rich elements (AREs) present in the 3’ untranslated region (UTR) of target mRNAs (Puig et al., 2005; Thompson et al., 1996). When Fe becomes scarce, Cth2 binds mRNAs that encode proteins involved in non-essential Fe-demanding pathways, including mitochondrial oxidative phosphorylation and TCA cycle, and promotes their degradation to prioritize Fe utilization to indispensable processes such as DNA synthesis and repair (Puig et al., 2005; Puig et al., 2008; Ramos-Alonso et al., 2020; Ramos-Alonso et al., 2018a). Disruption of the TZF RNA-binding domain or mutations of conserved cysteine (Cys) residues within the TZFs to arginine (Arg) prevent binding and decay of Cth2 target mRNAs (Puig et al., 2005; Vergara et al., 2011). More recently, Cth2 has been shown to directly inhibit the translation of several mRNA targets, in addition to its function in mRNA degradation (Ramos-Alonso et al., 2019; Ramos-Alonso et al., 2018a). However, the mechanisms of translational repression by Cth2 remain unknown.

Previous studies have documented dysregulation of Fe homeostasis and activation of the Fe regulon during the aging process. In yeast, impaired iron-sulfur cluster (ISC) synthesis has been shown to cause age-associated genomic instability (Diaz de la Loza Mdel et al., 2011; Veatch et al., 2009). Loss of vacuolar acidification is an early event during aging that has been linked to increased activity of the Fe regulon and mitochondrial dysfunction (Hughes and Gottschling, 2012). Consistent with these observations, deficiency of the vacuolar ATPase (V-ATPase), which is required for maintaining vacuolar acidity, shortens lifespan (Chen et al., 2020; Hughes et al., 2020; Schleit et al., 2013). In addition, a genetic screen has previously demonstrated that deletion of *CTH2,* which encodes for the Cth2 protein involved in post-transcriptional regulation of Fe homeostasis, as well as its close homolog *CTH1,* extends lifespan in yeast (McCormick et al., 2015).

While previous reports have shown that dysregulation of Fe homeostasis contributes to aging, the underlying molecular mechanisms are still not completely understood. In this study, we investigated how Fe homeostasis influences the aging process by performing unbiased analyses of the yeast deletion mutants lacking genes involved in Fe import and utilization. Here, we show that aging in yeast is associated with an up-regulation of Fe regulon genes whereas deletion of several members of the Fe regulon leads to increased longevity. We also show that constitutive activation of the Aft1 transcription factor shortens lifespan in *S. cerevisiae.* These findings demonstrate that expression of several Fe regulon genes is limiting lifespan in yeast. Finally, we reveal that deficiency of the RNA-binding protein Cth2 is sufficient to prevent the negative effect of Aft1 on lifespan by alleviating its repressive effects on its targets. Together, these findings uncover an important role of the RNA-binding protein Cth2 in the regulation of lifespan and suggest that modulation of Fe homeostasis may serve as an attractive strategy to delay aging and prevent the development of age-associated diseases in humans.

## Results

### Modulation of Fe metabolism impacts yeast replicative lifespan and cell fitness

To understand the impact of Fe homeostasis on cellular aging, we performed unbiased analysis of the effect of deleting genes involved in Fe import and utilization on lifespan in yeast *S. cerevisiae.* To this end, we analyzed replicative lifespan (RLS) of yeast strains lacking an Fe-responsive transcription factor (*AFT1*), as well as genes involved in cellular Fe uptake (*FET3, FIT2*), vacuolar Fe storage and mobilization (*CCC1, FTH1, VMA21*), mitochondrial Fe import (*MRS3*), iron-sulfur cluster (ISC) synthesis and distribution (*ISU1, GRX3*), recycling of heme iron (*HMX1*), and remodeling of cellular Fe metabolism (*CTH1, CTH2*) (Fig. 1A). Although most of the analyzed gene deletions involved in Fe homeostasis either did not significantly change lifespan or led to a shortened replicative lifespan, we found that deletion of *FET3, FIT2, HMX1, CTH1,* and *CTH2* genes significantly extended lifespan (p<0.05; Fig. 1B, Table S1). One of the longest-lived gene deletions, i.e., *CTH2* encoding an RNA-binding protein involved in Fe starvation response, extended lifespan by 51.1% (p<0.0001; Fig. 1B, Table S1). We next investigated how perturbations of Fe homeostasis impact cell fitness by analyzing the growth of the gene deletion mutants in the presence of different levels of Fe^2+^ in the media (Fig. S1A). We observed that the *aft1*Δ mutant has delayed growth on YPD medium. However, the cell fitness improved when the medium was supplemented with Fe in the form of ferrous ammonium sulfate (FAS). Consistent with prior reports, the growth of *aft1*Δ, *fet3*Δ, and *vma21*Δ cells was impaired on YPG medium containing 3% glycerol (which requires mitochondrial respiration to be utilized), indicating that these mutants are respiratory insufficient (Eide et al., 1993)) (Fig. S1B). However, supplementing YPG with FAS rescued the growth of *vma21*Δ cells, but not *aft1*Δ cells and only partially *fet3*Δ cells. In addition, Fe deficiency significantly delayed the growth rates of the *aft1Δ, fet3Δ, grx3*Δ and *vma21*Δ mutants (p<0.05) in the presence of Fe^2+^ chelator bathophenanthrolinedisulfonic acid (BPS) (Jo et al., 2009) (Fig. 1C). Together, these data suggest that the deletion of genes involved in different aspects of Fe homeostasis may differentially affect lifespan and cellular fitness.

**Fig. 1.**
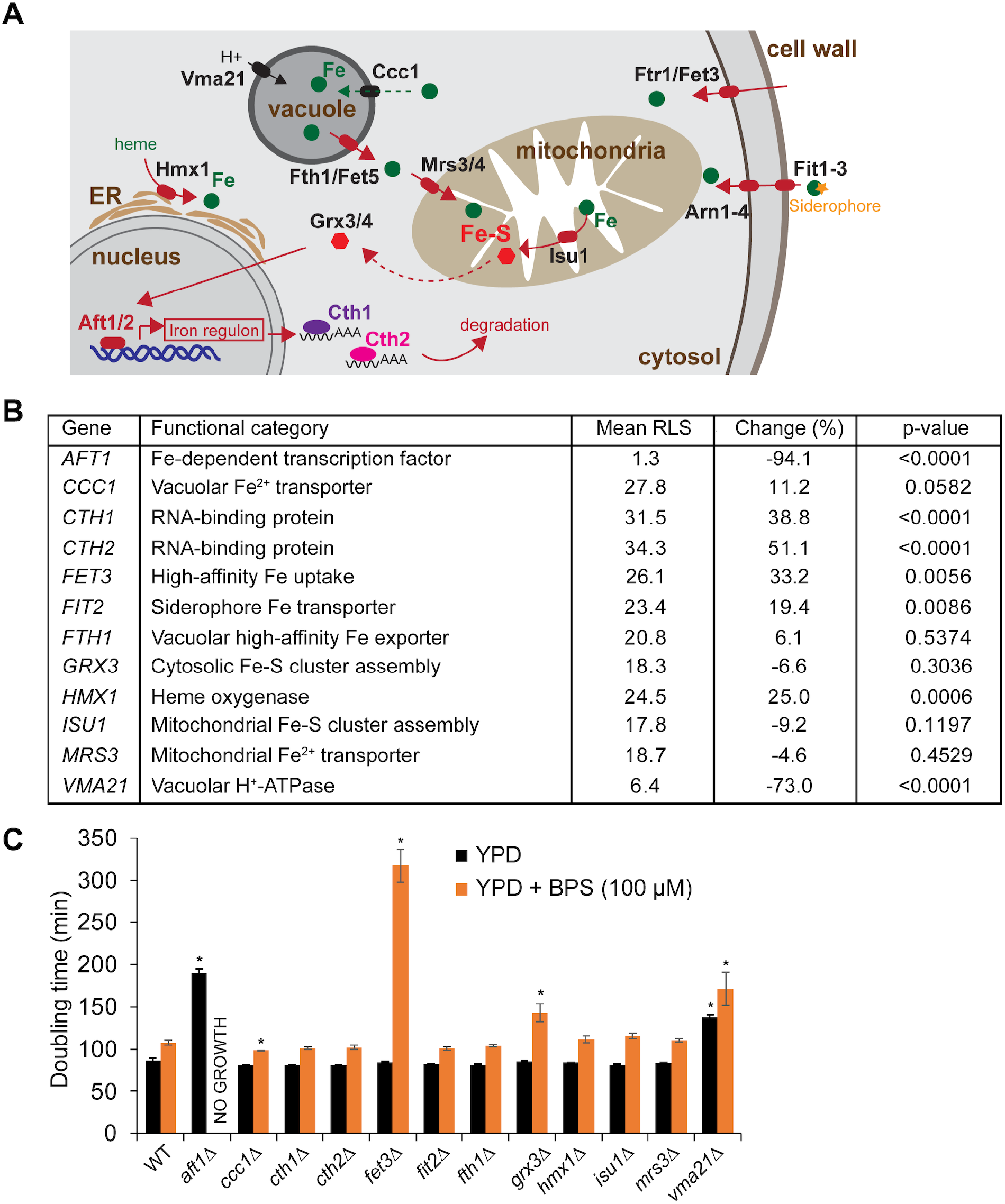
Deletion of genes involved in Fe homeostasis differentially affects lifespan and cell fitness. (**A**) Scheme of the Fe regulon in *S. cerevisiae.* Deficiency of the ISC production leads to nuclear localization of the Aft1 transcription factor, which activates expression of the Fe regulon genes involved in Fe assimilation. (**B**) Replicative lifespan of yeast mutants lacking genes involved in regulation of Fe homeostasis. (**C**) Doubling time in the absence or presence of 100 μM Fe^2+^ chelator BPS. Doubling time was calculated using the Yeast Outgrowth Data Analyzer (YODA) software. Error bars represent SEM (n=3). *p<0.05, compared to wild-type cells, Student’s t test.

### Old cells demonstrate a global decrease in mRNA translation, but an up-regulation of genes involved in Fe import into the cell

Previous studies have shown that protein synthesis is globally decreased in yeast during replicative aging (Hu et al., 2018). To determine whether overall inhibition of translation during aging leads to a decrease in expression of Fe-related proteins, we examined global protein synthesis levels in young and replicatively aged yeast using ribosome profiling (Ribo-Seq). For this, we isolated replicatively aged cells using the Mother Enrichment Program (MEP) (Hu et al., 2014; Lindstrom and Gottschling, 2009) combined with the biotin affinity purification step (Smeal et al., 1996). We isolated yeast cells that were cultured in the presence of estradiol for 2 hrs (YNG) and 30 hrs (OLD) and had undergone on average 2 and 15 cell divisions, respectively (Fig. 2A). Following enrichment of replicatively aged cells, mRNA expression and protein translation profiles were assessed in young and aged cells by RNA-Seq and Ribo-Seq approaches (Dataset S1). In order to account for differences in overall translational changes with aging, we spiked Ribo-Seq samples with 1% worm lysate at the beginning of the library preparation. The addition of the spike-in control from an evolutionarily distant organism allowed us to normalize the samples based on overall translation levels. By comparing the ratio of reads in each of the samples to the number of reads that align to the worm genome, we found that overall levels of protein translation were significantly decreased in replicatively aged cells compared to young cells (Fig. 2B).

**Fig. 2.**
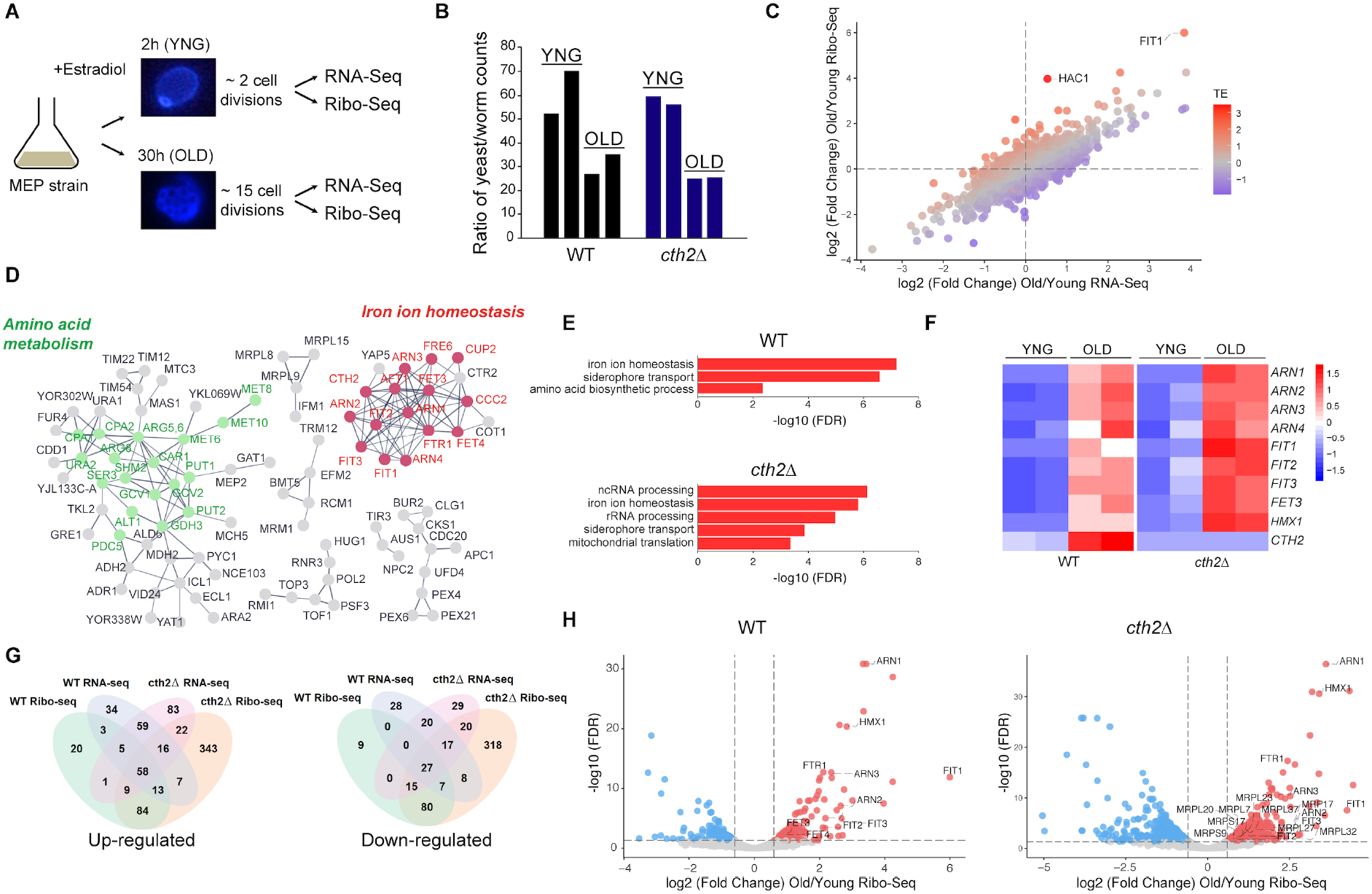
Aging leads to a global inhibition of translation, but an up-regulation of genes involved in Fe homeostasis. (**A**) RNA-Seq and Ribo-Seq analyses in young (YNG) and replicatively aged (OLD) cells isolated using the Mother Enrichment Program. (**B**) Spike-in enables direct comparison of footprints between Ribo-Seq samples. Two biological replicates are shown. (**C**) Comparison of transcriptional (RNA-Seq) and translational (Ribo-Seq) changes during aging. Genes whose translation efficiency (TE) is increased (red) or decreased (blue) with aging are highlighted. (**D**) Genes significantly up-regulated during aging (FDR<0.05) identified by Ribo-Seq were visualized using STRING (evidence view, high confidence). Genes without network partners were omitted. (**E**) Gene ontology analysis of the Ribo-Seq data indicates activation of pathways involved in Fe uptake during aging. (**F**) *CTH2* expression is upregulated during aging. Heatmap shows scaled expression values (normalized log2 Ribo-Seq read counts from Dataset S4) of the Fe regulon genes, including *ARN1, ARN2, ARN3, ARN4, FIT1, FIT2, FIT3, FET3, HMX1,* and *CTH2,* in young (YNG) and replicatively aged (OLD) wild-type and *cth2*Δ cells. (**G**) Venn diagrams show common significantly changed transcripts at the level of transcription (RNA-seq) and translation (Ribo-Seq) in wild-type and *cth2*Δ cells (FDR<0.05). (**H**) Cth2 is a negative regulator of mitochondrial translation. Volcano plot shows differentially translated genes (FDR > 0.05, Log2 fold change > 0.6) in wild-type and *cth2*Δ cells.

We next compared temporal changes in mRNA expression and protein translation in replicatively aged cells using the RNA-Seq and Ribo-Seq data (Dataset S2). Most of age-dependent changes detected by Ribo-seq correlated with the changes in the transcriptome (Fig. 2C) suggesting that most of the changes of gene expression during aging occur at the level of transcription. Despite the overall decrease in translation, our analysis revealed an up-regulation of genes involved in Fe ion homeostasis, siderophore transport, and amino acid biosynthesis with aging (Fig. 2D and 2E; Dataset S3). Interestingly, most of the up-regulated genes involved in Fe ion homeostasis are key components of the Fe regulon, a group of genes regulated by the Aft1 transcription factor (Martinez-Pastor et al., 2017). Together, our data indicate that aging is associated with a global inhibition of translation, but an up-regulation of genes involved in Fe import into the cell.

### Ribo-Seq analyses reveal mitochondrial translation as a novel post-transcriptional target of Cth2

Ribo-Seq analyses of replicatively aged cells showed that expression of the RNA-binding protein Cth2, which is involved in post-transcriptional repression of genes encoding for non-essential Fe-containing proteins, increases during aging (Fig. 2F). The fact that levels of Cth2 are increased with aging, whereas the deletion of *CTH2* extends lifespan suggests that Cth2 might be a negative regulator of longevity. To further understand the biological relevance of Cth2 during aging, we used Ribo-Seq to perform genome-wide analysis of translation in young and old *cth2*Δ mutant cells. Our analysis revealed that most changes observed in wild-type cells with aging were also detected in the *cth2*Δ mutant (Fig. 2G). Ribo-Seq identified 552 up-regulated genes and 492 down-regulated genes in the *cth2*Δ mutant with aging (FDR<0.05), of which 164 were commonly up-regulated with aging and 129 genes were commonly down-regulated with aging in both wild-type and the *cth2*Δ mutant. To distinguish between transcriptional and translational responses, we compared changes in gene expression in replicatively aged *cth2*Δ cells identified by the RNA-Seq and Ribo-Seq approaches. Although most of the changes in gene expression during aging were driven by changes in mRNA abundance, we identified 343 genes that were specifically up-regulated in the *cth2*Δ mutant at the level of translation (Fig. 2G).

Cth2 is an mRNA-binding protein that acts as a posttranscriptional regulator by affecting stability and translation of mRNAs containing the 5’-UAUUUAUU-3’ and 5’-UUAUUUAU-3’ AU-rich elements (AREs) in their 3’ untranslated (UTR) regions (Puig et al., 2005; Ramos-Alonso et al., 2018a). In response to Fe deprivation, Cth2 binds mRNAs encoding Fe-containing proteins leading to their degradation. In addition to known Cth2 targets, our data revealed that Cth2 is limiting the expression of genes involved in mitochondrial translation and genes encoding for mitochondrial ribosomal proteins. Among the genes translationally up-regulated in the *cth2*Δ mutant, we identified 29 genes involved in mitochondrial translation, including 10 ribosomal proteins of the small subunit and 16 ribosomal proteins of the large subunit (Fig. 2H and Fig. S2). However, most of these Cth2-dependent genes do not possess AREs in their 3’-UTR sequences suggesting that they might be coordinately regulated by Cth2, but not directly. Together, the evidence that Cth2 affects mitochondrial translation and the known role of Cth2 in limiting Fe for non-essential processes and down-regulating respiration, suggest Cth2 as a negative regulator of mitochondrial function during aging.

### Constitutive activation of the Fe regulon shortens lifespan in *S. cerevisiae*

We hypothesized that increased activity of Aft1 transcription factor in old cells leads to increased levels of Cth2, which might negatively regulate aging by repressing mitochondrial function. To test this hypothesis, we first asked whether constitutive activation of the Fe regulon in young cells is sufficient to shorten lifespan in *S. cerevisiae.* For this, we generated a yeast strain carrying a genome-integrated copy of the *AFT1-1^up^* allele (Yamaguchi-Iwai et al., 1995) and analyzed its replicative lifespan. This strain contains a mutation in the Aft1 protein that leads to its nuclear accumulation and constitutive expression of the Fe regulon genes, irrespective of the Fe availability. We found that constitutive activation of the Fe regulon by *AFT1-1^up^* allele significantly decreased the replicative lifespan of yeast cells (p<0.01; Fig. 3A, Table S1). To ascertain whether the up-regulation of *CTH2* by the *AFT1-1^up^* allele contributes to shortened lifespan in *AFT1-1^up^* expressing cells, we combined the constitutively active *AFT1-1^up^* allele with the *cth2*Δ deletion. Importantly, deletion of *CTH2* was able to rescue the short lifespan of *AFT1-1^up^* allele expressing cells (Fig. 3A) suggesting a negative effect of the increased Cth2 expression on lifespan.

**Fig. 3.**
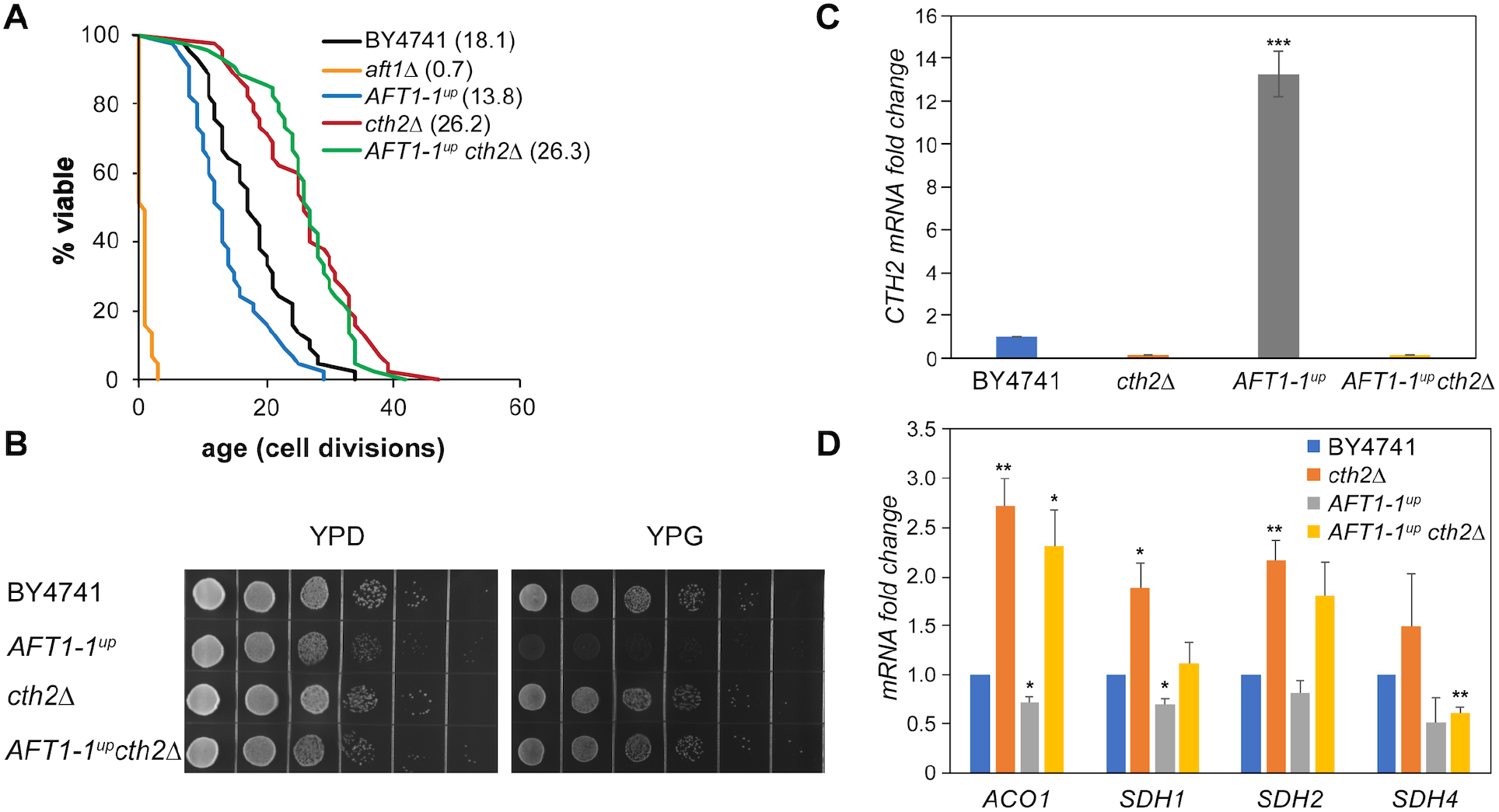
Constitutive activation of the Aft1 negatively regulates yeast lifespan. (**A**) Shortened lifespan in cells expressing *AFT1-1^up^* allele can be rescued by the deletion of *CTH2.* Mean lifespans are shown in parentheses. (**B**) Constitutive activation of the Aft1 transcription factor using *AFT1-1^up^* allele leads to inability to grow on glycerol containing medium (YPG), whereas deletion of *CTH2* is able to rescue this growth defect. 10× serial dilutions of logarithmically growing cells were spotted on agar plates with YPD (glucose) or YPG (glycerol) media. (**C**) Expression of *AFT1-1^up^* induces expression of Cth2. Relative mRNA levels of *CTH2* in exponentially growing cells were determined by RT-qPCR. Results are represented as means ± SEM of three independent experiments, ***p<0.001. (**D**) Deletion of *CTH2* alleviates its repressive effects on its targets. Relative mRNA levels of Cth2 targets in exponentially growing cells were determined by RT-qPCR. Results are represented as means ± SEM of three independent experiments, *p<0.05, **p<0.01.

Because many of the Cth2 targets include mRNAs encoding for mitochondrial proteins and components of the mitochondrial electron transport chain, one would expect that constitutive activation of Aft1 transcription factor would increase *CTH2* expression and inhibit mitochondrial function leading to the inability to grow on non-fermentable carbon sources that require respiration. The growth of the *AFT1-1^up^* allele expressing cells was significantly impaired on the non-fermentable medium containing glycerol as the only carbon source (Ramos-Alonso et al., 2018b) (Fig. 3B). Moreover, deletion of *CTH2* was able to rescue the growth of the *AFT1-1^up^* expressing cells on glycerol-containing medium. Finally, in agreement with its function in mRNA degradation, our data show that constitutive activation of the Aft1 transcription factor leads to increased expression of Cth2 (Fig. 3C) and Cth2-dependent post-transcriptional repression of nuclear-encoded components of the electron transport chain and the TCA cycle (Fig. 3D). These findings suggest that age-dependent activation of the Aft1 transcription factor could contribute to aging in *S. cerevisiae* and lead to impaired mitochondrial function by increasing expression of the Cth2 post-transcriptional regulator.

### Blocking mRNA-binding activity of Cth2 extends lifespan whereas increasing levels of Cth2 is sufficient to shorten lifespan in yeast

To investigate the underlying mechanisms by which Aft1-dependent activation of *CTH2* expression negatively regulates longevity, we asked whether blocking mRNA-binding activity of Cth2 is sufficient to extend lifespan. Cth2 contains an RNA-binding motif consisting of two tandem zinc fingers (TZFs), which directly interact with AREs within the 3’-UTR regions of its target mRNAs (Puig et al., 2005; Puig et al., 2008). Mutations of the conserved Cys residues within the TZF domain of Cth2 have been shown to abolish Cth2 binding of its target mRNAs and their subsequent degradation (Puig et al., 2005). To test whether the mRNA-binding function of Cth2 is directly related to its repressive effect on lifespan, we examined if mutation of the conserved Cys residue within the TZF domain to Arg (C190R) is sufficient to extend lifespan (Fig. 4A). We found that the replicative lifespan of the *CTH2-C190R* mutant was increased to a similar extent as the *cth2*Δ mutant (21.4%, p<0.01 compared to wild-type control; Fig. 4B, Table S1). These data demonstrate that the integrity of the TZFs and RNA-binding function of Cth2 are required for its repressive effects on lifespan.

**Fig. 4.**
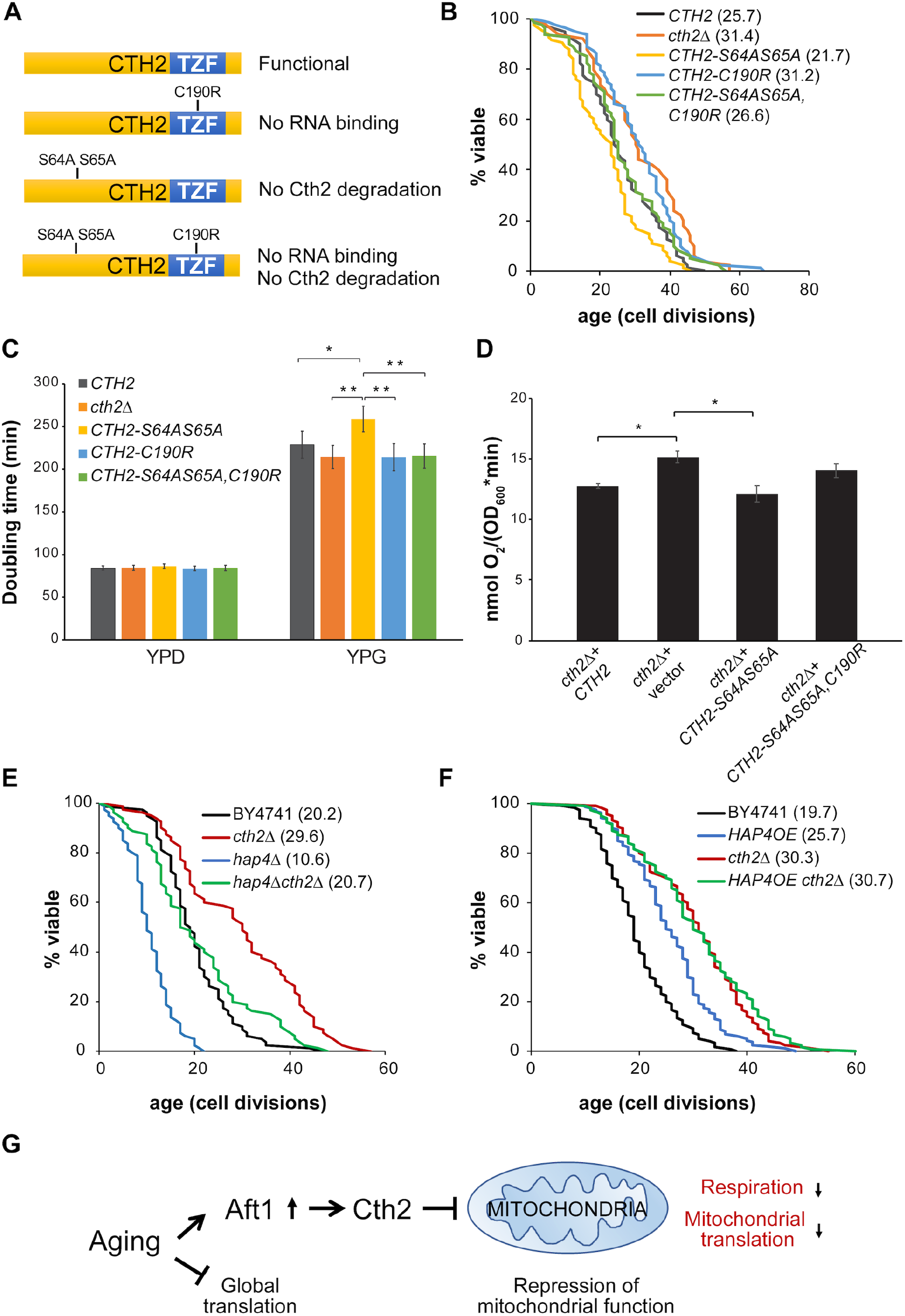
Cth2 is a negative regulator of lifespan and mitochondrial respiration. (**A**) Schematic representation of the Cth2 wild-type and mutant proteins used in this study. (**B**) Mutation of the conserved Cys residue within the TZF domain of Cth2 to Arg (C190R) is sufficient to extend lifespan. Mean lifespans are shown in parentheses. (**C**) Doubling times in YPD and the non-fermentable YPG media were calculated using Yeast Outgrowth Data Analysis (YODA) software. Error bars represent SEM (n=3), **p<0.01, *p<0.05. (**D**) Increased expression of Cth2 inhibits mitochondrial respiration. Oxygen consumption was measured in cells grown in YPG medium for 29 h. Error bars represent SEM (n=3), *p<0.05. (**E**) Extended lifespan in *cth2*Δ is dependent on the Hap4 transcription factor. (**F**) Hap4 overexpression (*HAP4OE*) and *cth2*Δ do not have an additive effect on lifespan. Mean lifespans are shown in parentheses. (**G**) Model showing the role of Cth2 in age-related repression of mitochondrial function. During aging, increased activity of the Aft1 transcription factor leads to increased expression of Cth2, which targets mRNAs involved in respiration and mitochondrial translation, leading to repression of mitochondrial function.

Recently, phosphorylation of a patch of serine residues within the N-terminal region of Cth2 has been shown to play an important role in its ubiquitin-dependent degradation (Romero et al., 2018). When Cth2 degradation is impaired by mutagenesis of the Cth2 serine residues, the levels of Cth2 protein increase leading to impaired growth in Fe-depleted conditions (Romero et al., 2018; Thompson et al., 1996). To further test how increased levels of Cth2 affect lifespan, we created a Cth2 double serine mutant (*CTH2-S64AS65A*) allele either alone or in combination with the TZF mutant (*CTH2-C190R)* (Fig. 4A). Our data demonstrate that the *CTH2-S64AS65A* (gain-of-function) mutant allele shortens lifespan (15.6%, p<0.05 compared to wild-type control; Fig. 4B, Table S1). Conversely, combining the Cth2 double serine mutant (*CTH2-S64AS65A*) with the TZF loss-of-function mutation (*CTH2-C190R*) abolished its negative effect on lifespan. Together, these data suggest that the non-degradable form of Cth2 protein has a deleterious effect on lifespan.

### Cth2 is a negative regulator of mitochondrial function

Previous studies revealed a correlation between oxygen consumption and increased levels of mitochondrial respiration with lifespan extension in yeast (Barros et al., 2004; Bonawitz et al., 2007; Lin et al., 2002). Because targets of Cth2 include components of the mitochondrial electron transport chain and TCA cycle, we hypothesized that increased levels of Cth2 would inhibit mitochondrial function leading to decreased respiration. To test this, we analyzed the growth of cells expressing the non-degradable double serine Cth2 mutant (*CTH2-S64AS65A*), the TZF domain mutant (*CTH2-C190R*), and the *CTH2-S64AS65A,C190R* mutant in YPG medium containing glycerol as carbon source that requires respiration (Fig. 4C). We observed that the population doubling time of the cells expressing a non-degradable form of Cth2 (*CTH2-S64AS65A*) was significantly increased (p<0.01) in YPG medium compared to *cth2*Δ and *CTH2-C190R* mutant cells. Moreover, introducing the C190R mutation that blocks RNA-binding activity of Cth2 completely prevented this effect. To further assess that this effect is due to decreased respiration, we measured the levels of oxygen consumption in YPG medium in the *cth2*Δ mutant as well as in cells expressing *CTH2-S64AS65A* and *CTH2-S64AS65A,C190R* mutations (Fig. 4D). We found that *CTH2* loss of function (*cth2*Δ) improves respiration compared to cells expressing wild-type *CTH2*. In turn, expression of the *CTH2-S64AS65A* mutation significantly decreased (p<0.05) oxygen consumption, whereas mutation in the TZF domain (*CTH2-S64AS65A,C190R*) abolished this effect.

Overexpression of the Hap4 transcription factor, which controls the expression of many genes encoding components of the mitochondrial electron transport chain, has been shown to extend *S. cerevisiae* lifespan by increasing respiration (Lin et al., 2002). Given that Cth2 affects the expression of the respiratory genes at the post-transcriptional level, we asked whether increased lifespan in the *cth2*Δ mutant is dependent on the Hap4 transcription factor. Consistent with this possibility, deletion of *HAP4* significantly decreased the lifespan in the *cth2*Δ background (30.1%, p<0.001; Fig. 4E, Table S1). To further validate that Cth2 deficiency extends lifespan by affecting mitochondrial respiration, we performed an epistasis experiment combining *cth2*Δ with Hap4 overexpression (*HAP4OE).* As expected, combining *cth2*Δ with *HAP4OE* had no additive effect on lifespan (Fig. 4F, Table S1). Taken together, our data suggest that Cth2 is a negative regulator of lifespan, which accumulates with aging and down-regulates its downstream targets leading to Cth2-dependent repression of mitochondrial respiration.

## Discussion

Dysregulation of Fe homeostasis has been implicated in the development of many human age-related diseases such as type 2 diabetes, cancer, neurodegeneration, and cardiovascular disease. However, how Fe homeostasis contributes to organismal aging remains unclear. Previous studies have also shown that aging in model organisms is associated with a decline of mitochondrial function and activation of genes involved in Fe homeostasis, but the link between these processes and aging is not completely understood (Sun et al., 2016; Veatch et al., 2009). Here, we performed an unbiased screen to investigate how deletion of individual genes involved in Fe import and utilization affects lifespan using yeast *S. cerevisiae* as a model. Although the majority of the tested deletion mutants did not significantly change lifespan, we found that deletion of several members of the Fe regulon, including *FET3, FIT2, HMX1, CTH1* and *CTH2,* led to increased longevity (Fig. 1). These data, together with the observation that activity of the Fe regulon is increased during aging, suggest a model that increased expression of the Aft1 targets might limit replicative lifespan in yeast. In support of this model, our data show that expression of the constitutively active Aft1 mutant (*AFT1-1^up^*) allele, which activates Fe regulon genes irrespective of the Fe availability, leads to shortened lifespan.

One of the longest-lived mutants we analyzed lacks *CTH2,* a gene encoding an RNA-binding protein that post-transcriptionally regulates Fe-related proteins during Fe deficiency (McCormick et al., 2015). *CTH2* is also one of the major transcriptional targets that are activated by the Aft1 transcription factor, which showed a robust increase during aging. Here, we propose that Cth2-dependent repression of its targets mediates the negative effects of Aft1 on lifespan leading to repression of mitochondrial function. In agreement with this hypothesis, deletion of *CTH2* in cells expressing *AFT1-1^up^* allele was able to abolish the negative effect of *AFT1-1^up^* on lifespan. Since Cth2 targets include many mitochondrial genes involved in TCA cycle and respiration (Puig et al., 2005; Ramos-Alonso et al., 2018a), one would expect that increased expression of Cth2 would inhibit mitochondrial function leading to decreased respiration. Indeed, our data demonstrate that Cth2 negatively regulates respiration. In addition, we show that the *AFT1-1^up^* allele expressing cells were unable to grow on non-fermentable carbon sources, but deletion of *CTH2* was sufficient to alleviate the negative effects of Aft1 expression on respiration.

Due to its involvement in the regulation of respiration and other Fe-dependent pathways, derepression of Cth2 targets in the *cth2*Δ mutant may explain the effect of Cth2 loss on lifespan. Mutation of the conserved Cys residue within the TZF domain of Cth2 that inhibits its mRNA-binding activity was sufficient to confer longevity, whereas Cth2 gain-of-function shortened replicative lifespan. These data suggest that deletion of *CTH2* extends lifespan by directly alleviating mRNA-binding and its repressive effects on its targets. Specifically, we show that that expression of constitutively active *AFT1-1^up^* allele leads to increased expression of Cth2 and Cth2-dependent repression of the components of the electron transport chain and the TCA cycle leading to repression of mitochondrial function. Consistent with the repressive effect of Cth2 on respiration, deletion of *GRR1* gene in yeast has been shown to prevent the growth of cells on media with non-fermentable carbon sources (Romero et al., 2018). Grr1 is the protein that recognizes phosphorylated Cth2 and facilitates its proteasomal degradation (Mark et al., 2014; Romero et al., 2018). Therefore, elevated Cth2 protein levels in the *grr1*Δ mutant might inhibit the growth of cells on non-fermentable carbon sources through Cth2-mediated degradation of its targets. The role of Cth2 as a negative regulator of mitochondrial function is further supported by the fact that deletion of *CTH2* in *grr1*Δ prevented the negative effect of Cth2 on growth in YPEG medium (Romero et al., 2018). Moreover, we demonstrate that overexpression of the Hap4 transcription factor, which has been shown to extend lifespan by increasing respiration (Li et al., 2020; Lin et al., 2002), had no additive effect on lifespan with the deletion of *CTH2.* Together, these data suggest that Cth2 is a negative regulator of mitochondrial function and longevity.

We also identified mitochondrial translation as a novel post-transcriptional target of Cth2. Although many of the prior studies investigated changes in gene expression during aging at the transcriptional level across multiple species, the role of post-transcriptional regulation in aging is much less understood. By applying a combination of the RNA-Seq and Ribo-Seq approaches, we identified genes that are specifically regulated by Cth2 at the post-transcriptional level during aging. Although many of the mitochondrial ribosomal proteins were activated in the *cth2*Δ mutant with aging, most of them do not contain putative ARE sequences in their 3’-UTR. One possibility is that these proteins are regulated by a common transcription or translation factor that is directly regulated by Cth2. Further studies are required in order to investigate the mechanism of how Cth2 regulates mitochondrial translation. Together, Cth2-dependent inhibition of mitochondrial respiration and repression of mitochondrial translation may contribute to mitochondrial dysfunction during aging.

In summary, our data demonstrate that dysregulation of Fe homeostasis negatively regulates longevity through the activity of the Aft1 transcription factor and the Cth2 mRNA-binding protein function (Fig. 4G). Notably, the effects of Cth2 on regulation of lifespan are evolutionarily conserved, as deficiency of the orthologous protein in *Caenorhabditis elegans,* pos-1, significantly extends the animal’s lifespan (Curran and Ruvkun, 2007; Smith et al., 2008). Given that the mammalian homolog of Cth2, called tristetraprolin (TTP), also post-transcriptionally modulates the expression of Fe metabolism-related genes including transferrin receptor 1 and genes involved in mitochondrial electron transport chain (Bayeva et al., 2012; Sato et al., 2018), our data open a possibility that modulation of Fe starvation signaling can serve as a target for potential aging interventions in humans.

## Materials and Methods

### Yeast Strains and Plasmids

The yeast strains used in this study and their genotypes are listed in Table S2. Oligonucleotides used for gene disruption and cloning are listed in Table S3. Cells were grown at 30°C in standard YPD medium (1.0% yeast extract, 2.0% peptone, and 2.0% glucose) unless otherwise stated. One-step PCR-mediated gene disruption was performed using standard techniques. All genotypes were verified using single colony screening with polymerase chain reaction (PCR). The yeast Mother Enrichment Program (MEP) strain (UCC8773) was used to generate old and young cell ribosome profiling libraries (Henderson et al., 2014; Lindstrom and Gottschling, 2009). MEP cells were grown under nourseothricin (NAT) (100 μg/mL) and Hygromycin B (300 μg/mL) selection.

To create yeast strain BY4741 with *CTH2, CTH2*-C190R, *CTH2*-S64AS65A, and *CTH2*-S64AS65A,C190R, their corresponding ORFs were amplified from plasmids pSP414, pSP429, pSP853, and pSP898, respectively (Romero et al., 2018). PCR amplified *CTH2* and *CTH2* with mutations were then cloned into pDZ415 by using NotI and SalI restriction sites upstream of loxP-KanMX-loxP. Then *CTH2* and *CTH2* with mutations along with loxP-KanMX-loxP were amplified using the primers flanked on the 5’ side by sequence homologous to *CTH2* ORF and the 3’ side by sequence homologous to *CTH2* 3’-UTR for genomic integration. After the integrations were confirmed with colony PCR, cells were transformed with 0.2 μg pSH47 (Cre recombinase) for removal of KanMX marker and the transformants were selected on SC lacking uracil. After the transformation of the pSH47, cells were incubated overnight in YP medium supplemented with 3% galactose for the induction of Cre recombinase. The culture was subsequently diluted in 10fold dilutions and plated out on YPD medium and incubated at 30 °C for 30 h. The resulting colonies were replica-plated on YPD and YPD with 200 μg/ml G418 for the selection of colonies that lost the KanMX marker and verified by colony PCR. *AFT1-1^up^* sequence was amplified using pAFT1-1UP (Ueta et al., 2012) and similar approach that was used to create *CTH2* mutant yeast strains was used to generate yeast strain with genomically expressing *AFT1-1^up^* allele.

To create *ADH1* promoter driven *HAP4,* a 705 bp fragment of *ADH1* promoter was cloned into pRS306 using NotI and XhoI restriction sites. Then URA3 marker along with *ADH1* promoter was PCR amplified using the primers flanking the 5’ end by sequence homologous to *HAP4* promoter and the 3’ end by sequence homologous to *HAP4* ORF for genomic integration.

### Spot Assays

The growth of strains on in YPD and YPG media containing 3% glycerol as carbon source was determined using spot assays. Cells were initially grown in liquid culture until OD_600_ = 0.6, and 10× serial dilutions for each strain were spotted on YPD or YPG agar plates in the absence and presence of 0.5 mM ferrous ammonium sulfate. The plates were incubated at 30°C, and images were taken 48 h after plating.

### Replicative Lifespan

Lifespan assays were carried out as described (Steffen et al., 2009). Cells were grown on freshly prepared YPD plates at 30°C. Cells were monitored for cell divisions until cells stopped dividing and subsequent daughter cells were removed using a micromanipulator. Replicative lifespan was calculated as the number of times each mother cell divided before it underwent senescence.

### Growth Rate Analyses

Yeast growth rates were analyzed using the Epoch 2 Microplate Spectrophotometer (BioTek) and the doubling time interval was calculated using the yeast outgrowth data analyzer (YODA) as described (Lee et al., 2017; Olsen et al., 2010). Results are represented as means ± SEM of at least three independent experiments. Statistical significance of the data was determined by calculating p values using Student’s t test.

### Isolation of young and old yeast cells

Single colonies were inoculated in 5 mL YPD containing NAT and Hygromycin B and grown overnight at 30°C. The next day cells were diluted to OD_600_=0.2 into 20 mL of the fresh medium and grown until cells reach log phase. Cells were collected by centrifugation at 3,000 rpm for 3 min, washed twice in sterile PBS and 300×10^6^ cells were used for 30 min biotin labeling (5 mg/mL, EZ-Link Sulfo-NHS-LC-Biotin, Thermo Fisher). Excess Biotin was quenched with 0.1 M Glycine and cells were resuspended in sterile PBS before inoculation into fresh YPD medium containing 1 μM estradiol to initiate the Mother Enrichment Program. The 300×10^6^ cells were split between young (2 hrs after induction) and old (30 hrs after induction). Cells were collected by centrifugation at 3,000 rpm for 3 min, resuspended in ice cold PBS+BE (1 mg/mL BSA + 2 mM EDTA) and incubated for an hour at +4°C with Dynabeads Biotin Binder (Thermo Fisher). Cells were repeatedly washed with ice-cold sterile PBS while on magnet. Finally, sorted cells were counted and recovered in regular YPD 2% glucose for 30 min at 30°C on shaker before collection, flash frozen in liquid nitrogen and stored at −80°C. For each condition, two cell sortings were combined in order to reach at least 100×10^6^ cells necessary to make the RNA-Seq and ribosome profiling libraries.

### Ribosome Profiling

Yeast extracts were prepared by cryogrinding the cell paste with BioSpec cryomill. The cell paste was re-suspended in lysis buffer (20mM Tris-HCl pH8, 140mM KCl, 5mM MgCl, 0.5mM DTT, 1% Triton X-100, 100 μg/mL cycloheximide), spun at 20,000xg at 4°C for 5 min and 1 mL of the supernatant was transferred to a new Eppendorf tube. To normalize raw reads obtained from young and old cells, an equal amount of worm lysate was added to final concentration of 1% to each sample. After homogenization and spike-in normalization, samples were divided into two tubes, for total mRNAs and footprint extraction. The RNA-seq and ribosome profiling libraries were prepared as described (Beaupere et al., 2017) and sequenced using the Illumina HiSeq platform. Ribosomal footprint and total mRNAs reads were aligned to the *S. cerevisiae* genome from the Saccharomyces Genome Database (https://www.yeastgenome.org/, release number R64-2-1) and to the *C. elegans* genome (NCBI, WBcel235 assembly, RefSeq GCF_000002985.6) for linear normalization (Chen et al., 2015; Hu et al., 2018). Sequence alignment was performed using STAR software 2.7.1a, allowing two mismatches per read (Dobin et al., 2013). Counts were generated with featureCounts from Rsubread 1.22.2 package (Liao et al., 2019). Identification of differentially expressed genes was performed using generalized linear model of the EdgeR package, FDR<0.05 (Robinson et al., 2010).

### RT-qPCR

Total RNA was isolated by hot acid phenol extraction. RNA was treated with DNaseI, and 1 μg of RNA was used for cDNA synthesis using SuperScript III reverse transcriptase (Thermo Fisher Scientific) with random hexamer primers according to manufacturer’s instructions. mRNA expression was then analyzed by real-time PCR using KAPA SYBR FAST qPCR Master Mix (Kapa Biosystems) and the CFX-96 Touch Real-Time PCR Detection System (Bio-Rad Laboratories). *ACT1* was used as a reference gene for normalization of mRNA expression between genotypes. Results are represented as means ± SEM from three independent experiments.

### Analysis of oxygen consumption

Oxygen uptake by cells was measured after growing cells in YPG medium for 29 h using a Clark-type oxygen electrode (Oxyview system, Hansatech). The oxygen consumption rate was expressed as nmol O2 consumed per minute per OD_600_ (nmol O_2_/(min X OD_600_)) and was taken as an index of the respiratory ability.

### Statistical analysis

Statistical significance of the RT-qPCR data was determined by calculating p values using Student’s t test. Error bars represent standard errors of the means (SEM) unless otherwise noted. The summary and statistics of the lifespan assays are shown in Table S1. Statistical significance of the lifespan data was determined using the Wilcoxon Rank-Sum test (Wilcoxon, 1946).

## Data Availability

Raw reads and processed sequencing data generated by this study have been deposited in the GEO database (www.ncbi.nlm.nih.gov/geo/; Accession: GSE189306).

## Acknowledgments

We would like to thank Dr. Jerry Kaplan for providing the pAFT1-1UP plasmid used in this study. This work was supported by the NIH Grants AG058713 and AG066704 (to V.M.L.).

