## Supplementary Information for "Deficiency of the RNA-binding protein Cth2 extends yeast replicative lifespan by alleviating its repressive effects on mitochondrial function"

### Supplementary Figures

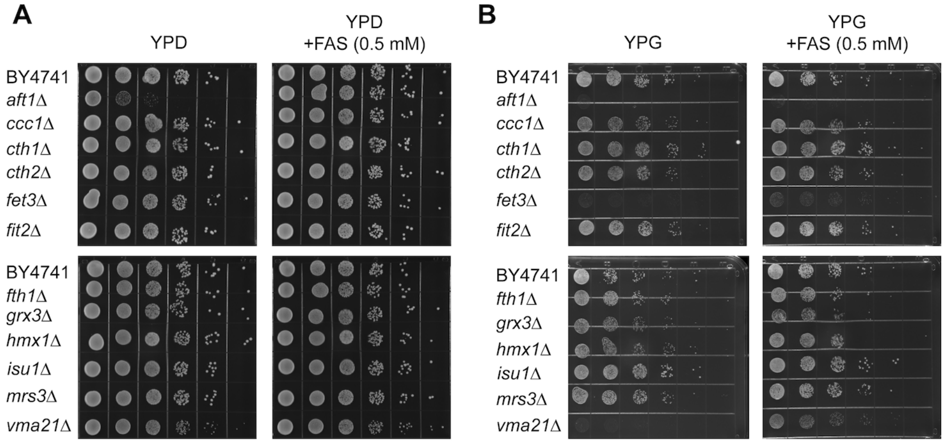


**Fig. S1.** Deletion of genes involved in Fe metabolism differentially affects cell fitness. (**A**) Growth of the gene deletion mutants on rich YPD media in the absence and presence of 0.5 mM ferrous ammonium sulfate (FAS). (**B**) Growth of the mutants on non-fermentable YPG media containing 3% glycerol as a carbon source in the absence and presence of 0.5 mM ferrous ammonium sulfate (FAS). 10× serial dilutions of logarithmically growing cells were spotted and incubated for 48 h at 30°C.

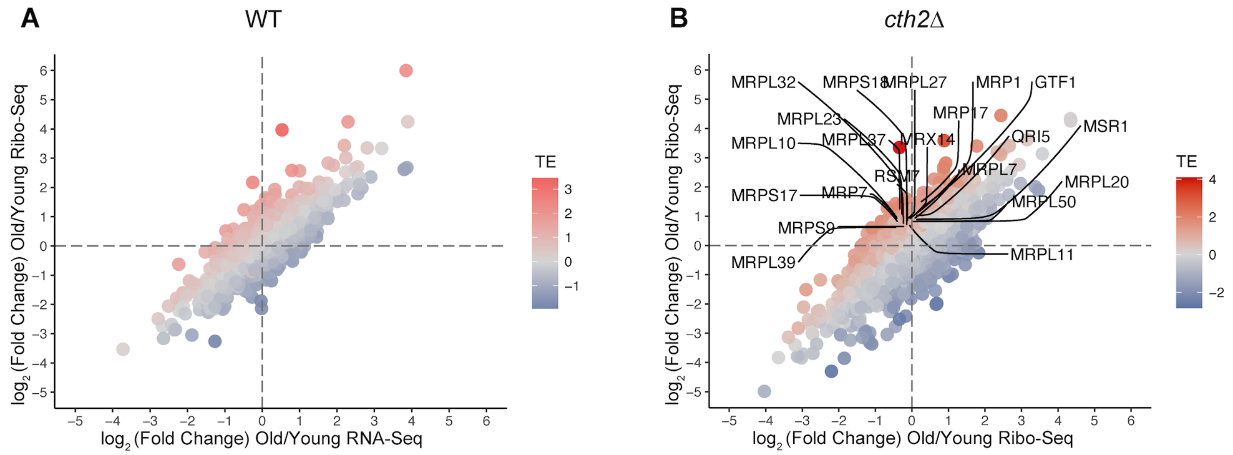


**Fig. S2.** Comparison of transcriptional (RNA-Seq) and translational (Ribo-Seq) changes during aging in wild-type (**A**) and *cth2Δ* (**B**) cells. Log2 changes in translation efficiency (TE) during aging were calculated with EdgeR. Genes involved in mitochondrial translation whose Log2 fold change in TE is greater than 0.6 are shown.
